# In Vivo Sublayer Analysis Of Human Retinal Inner Plexiform Layer Obtained By Visible-Light Optical Coherence Tomography

**DOI:** 10.1101/2021.01.08.425925

**Authors:** Zeinab Ghassabi, Roman V. Kuranov, Mengfei Wu, Behnam Tayebi, Yuanbo Wang, Ian Rubinoff, Xiaorong Liu, Gadi Wollstein, Joel S. Schuman, Hao F. Zhang, Hiroshi Ishikawa

## Abstract

**Purpose:** Growing evidence suggests, in glaucoma, the dendritic degeneration of subpopulation of the retinal ganglion cells (RGCs) may precede RGCs soma death. Since different RGCs synapse in different IPL sublayers, visualization of the lamellar structure of the IPL could enable both clinical and fundamental advances in glaucoma understanding and management. In this pilot study, we investigated whether visible-light optical coherence tomography (vis-OCT) could detect the difference in the inner plexiform layer (IPL) sublayers thicknesses between small cohorts of healthy and glaucomatous subjects.

**Method:** We investigated vis-OCT retinal images from nine healthy and five glaucomatous subjects. Four of the healthy subjects were scanned three times each in two separate visits, and five healthy and five glaucoma subjects were scanned three times during a single visit. Raster speckle-reduction scans (3 by 3 by 1.2 mm^3: horizontal; vertical; axial directions with 8192×8×1024 samplings, respectively) of the superior macular were acquired. IPL sublayers were then manually segmented using averaged A-line profiles.

**Results:** The mean ages of glaucoma and healthy subjects are 59.6 +/- 13.4 and 45.4 +/- 14.4 years (p =0.02, Wilcoxon rank-sum test), respectively. The visual field mean deviation (MD) are −26.4 to −7.7 dB in glaucoma patient and −1.6 to 1.1 dB in healthy subjects (p =0.002). The mean circumpapillary retinal nerve fiber layer (RNFL) thicknesses are 59.6 +/- 9.1 μm in glaucoma and 99.2 +/- 16.2 μm in healthy subjects (p=0.004). Median coefficients of variation (CVs) of intra-session repeatability for the entire IPL and three sublayers are 3.1%, 5.6%, 6.9%, and 5.6% in healthy subjects and 1.8%, 6.0%, 7.7%, and 6.2% in glaucoma patients, respectively. The mean entire IPL thicknesses are 36.2 +/- 1.5 μm in glaucomatous and 40.1 +/- 1.7 micrometer in healthy eyes (p=0.003, Mixed-effects model). We found that the middle sublayer thickness was responsible for the majority of the difference (14.2 +/- 1.8 μm in glaucomatous and 17.5 +/- 1.4 in healthy eyes, p<0.01).

**Conclusions:** IPL sublayer analysis revealed that the middle sublayer could be responsible for the majority of IPL thinning in glaucoma. Vis-OCT quantified IPL sublayers with good repeatability in both glaucoma and healthy subjects. Visualization of the IPL sublayers may enable the investigation of lamella-specific changes in the IPL in glaucoma and may help elucidate the response of different types of RGCs to the disease.

## Introduction

Glaucoma is a neurodegenerative disease characterized by retinal ganglion cell (RGC) death and axon degeneration, leading to vision loss ^1, 2^. Over the past two decades, optical coherence tomography (OCT) provided repeatable *in vivo* quantitative thickness assessment of retinal nerve fiber layer (RNFL) and combined ganglion cell layer (GCL) and inner plexiform layer (IPL) referenced as GCIPL, which are clinically useful biomarkers for glaucoma assessment ^3–7^

RGCs have complex yet characteristic dendritic morphology that determines how they receive and transmit visual information^8^. Specifically, the inner neurons form synapses with RGC dendrites in the IPL, which can be divided into ON and OFF sublamellae, reflecting the functional segregation of the ON and OFF pathways^8, 9^ ON RGCs have dendritic arbors in the inner region of the IPL, while OFF RGC dendrites co-localize with OFF bipolar axonal terminals in sublamella *a*^10, 11^. ON-OFF RGCs have dendrites arborizing in both sublamellae *a* and *b* of the IPL and respond to both light onset and offset ^12^ Since IPL consists of various types of dendrites, a quantitative analysis of the IPL sublayer structure may provide additional information about glaucomatous insults to the retinal neural tissues *in vivo*.

However, the changes of the ON and OFF IPL sublamellae in glaucoma is controversial. It was shown in *ex-vivo* studies in mice that the IPL layer can be the first location of the structural glaucomatous damage^13^. Although some studies indicate that an OFF sublamella is affected in glaucoma ^14, 15^, while other studies suggested that an ON sublamella was susceptible to the optic nerve crush (ONC) ^16, 17^.

Recently, the new OCT modality that utilizes the visible light spectrum (vis-OCT), in contrast to the conventional near-infrared (NIR) spectrum, made it possible to delineate finer retinal layers ^18–24^, which is not feasible using NIR OCT ^25–27^ Such an improvement in retinal layer imaging is due to the increased scattering contrast and the axial resolution at shorter wavelengths. Although NIR ultra-high-resolution (UHR) OCT axial resolution of 1.8 μm ^27^ is approaching the vis-OCT resolution of 1.3 μm used in our study, the higher resolution alone is not sufficient for robust visualization of the sublayer laminations, such as IPL and outer retina. For improved visualization and quantification of the fine retinal layers, higher contrast and mitigation of the speckle blurring effect are required. The shorter wavelength using in vis-OCT provides such required contrast. And the recently reported speckle reduction techniques ^28^ in vis-OCT ^22, 29^ mitigated the speckle blurring effect in this work.

Our vis-OCT study revealed three hyper-reflective and two hypo-reflective bands in the IPL ^30^, corresponding well with the standard anatomical division of the IPL into five strata ^7^. Among those five IPL bands, we identified three IPL sublayers using the minimal vis-OCT signal intensity of the two hypo-reflective bands and the outer IPL boundaries. We measured IPL sublayer thickness in a small cohort of glaucoma and healthy subjects to evaluate vis-OCT IPL sublayer imaging repeatability and potentially quantify dendritic degeneration of the RGCs in glaucoma.

## Methods

### Subjects recruiting

The study was approved by the New York University Langone Health institutional review board and complied with the tenets of the Declaration of Helsinki. Informed consent was received from all subjects before imaging. Both men and women of all races/ethnicities ages 18 years or older were eligible for the study.

Fourteen eyes of 14 subjects (nine healthy and five glaucomatous) were enrolled. Five healthy and five glaucoma subjects participated in the intra-session repeatability study, while four healthy subjects also participated in the inter-session repeatability study. For the intra-session repeatability study, both glaucoma and healthy subjects were imaged three times in a single visit. For inter-session repeatability, four healthy subjects were imaged three times each in two separate visits.

All subjects were tested with visual field (VF), commercial NIR OCT, and vis-OCT. VFs were tested with the Swedish interactive thresholding algorithm 24-2 perimetry (SITA standard; Humphrey Field Analyzer; Zeiss, Dublin, CA). Reliable VFs were considered tests with less than 33% fixation losses and false-positive and false-negative responses. Mean deviation (MD) was used for the analysis. All subjects were imaged with NIR OCT (Cirrus HD-OCT; Zeiss) using the optic nerve head (ONH) cube of 200 × 200 scans. Global mean circumpapillary RNFL thickness was used for analysis.

To be eligible for the study, healthy individuals must exhibit the following characteristics: IOP < 21 mmHg; normal VF - glaucoma hemifield test (GHT) within normal limits, where one test is sufficient given acceptable reliability; normal-appearing ONH; intact neuroretinal rim without splinter hemorrhages, notches, or localized pallor or RNFL defect; symmetric ONH between eyes; and cup-to-disc ratio (CDR) difference < 0.2 in both vertical and horizontal dimensions.

To be eligible for the study, glaucoma individuals must exhibit both structural and functional vision loss as indicated by the following parameters: glaucomatous VF defect - GHT outside of normal limits; at least two tests needed within seven month period; and abnormal NIR OCT (mean RNFL < 70 microns).

### Vis-OCT imaging

We used the Aurora X1 vis-OCT system (Opticent Inc., Evanston, IL) ^21, 22^ to image IPL sublayers. The system was running at a 25,000 A-lines/sec rate. The incident power was set below 250 μW, which was within the laser safety limit defined by the ANSI standard ^31, 32^ Vis-OCT irradiation power was measured using a calibrated power meter (PM100D, Thorlabs, NJ) before each imaging session.

We used a unique speckle-reduction raster scanning protocol for vis-OCT image acquisition ^21, 22^ The scanning covers a volume of 3×3×1.2 mm^3^ (horizontal×vertical×axial with 8192×8×1024 pixels, respectively) in the retina centered at the foveola along the x-axis (Figure 1a). Along the y-axis, the data set was acquired from the superior part of the fovea, where one B-scan (the last or second last B-scan) crossed the foveola. Along the x-axis, we acquired a speckle-reduced B-Scan (srB-scan) image by vertically translating the vis-OCT scanning beam along the y-axis, as shown in Figure 1b. Such a spatial translation consisted of eight uncorrelated A-lines, as highlighted by the red dots in Figure 1b, with an interval of 6 μm, and the spatial interval between two vertical translation was 2.9 μm, as shown in Figure 1b. For every two vertical translations, we calculated a speckle-reduced A-line (srA-line) by averaging 16 regular A-lines as highlighted by the green dashed box in Figure 1b. Therefore, each srB-scan contains 512 srA-lines, and the spatial interval between the srB-scans is 375 μm.

**Figure 1.**
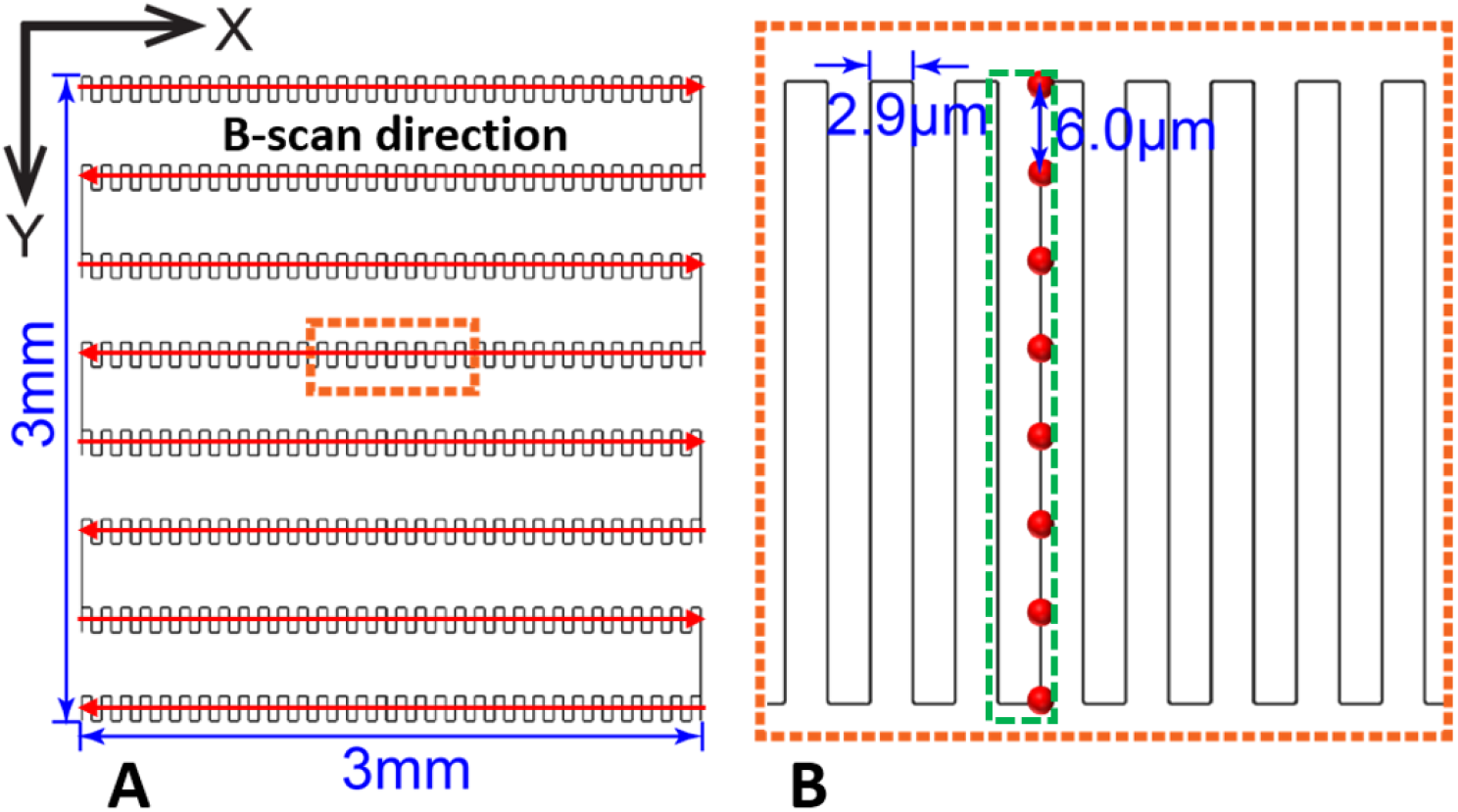
(a) Illustration of the overall speckle-reduction raster scanning protocol in vis-OCT; (b) Illustration of srA-line acquisition as highlighted by the dashed box in panel a. The red dots indicate the spatial locations of each regular A-line.

### Post-processing and measurement sampling

Figure 2 illustrates the method of calculating IPL layer thickness and its underlying anatomical lamination. The initial reference data cube was acquired, and an srB-scan crossing the foveola was identified. Next, a quality index (QI) ^33^ was computed for all srB-scans of the reference data cube that is superior to the srB-scan crossing the foveola, and a reference srB-scan (Figure 2a) with the highest QI was selected to measure the IPL sublayer thicknesses. In the following imaging sessions, we identified srB-scan from the same location from the current data cube by pinpointing a blood vessel pattern specific to the reference sr-Bscan (e.g., vessels 1, 2, and 3 in Figure 2a). All measurements were performed at the same distance from the selected vessel in the pattern in all scans of the subject. We then identified a segment of the IPL layer consisted of 15 srA-lines (Figure 2b) to obtain a depth-resolved OCT amplitude (tissue reflectivity) profile (averaged A-line; a blue curve in Figure 2c) to reduce noise in actual thickness variation measurements. In the averaged A-line, three peaks and two valleys can be identified (Figure 2c) corresponding to high- and low-intensity bands in the srB-scans (Figures 2a and 2b). Such sublamination revealed in the srB-scan was well correlated with the anatomical five strata of the IPL reported in the literature ^8–12^ The hypothesized correspondence of the five strata (S1-S5) with measured IPL sublayers (L1-L3) was indicated in Figures 2c and 2d. A simplified sketch explaining the origin of the five strata as a stratification of dendrites from ON and OFF center ganglion cells and bi-laminating ganglion cells is shown in Figure 2d.

**Figure 2.**
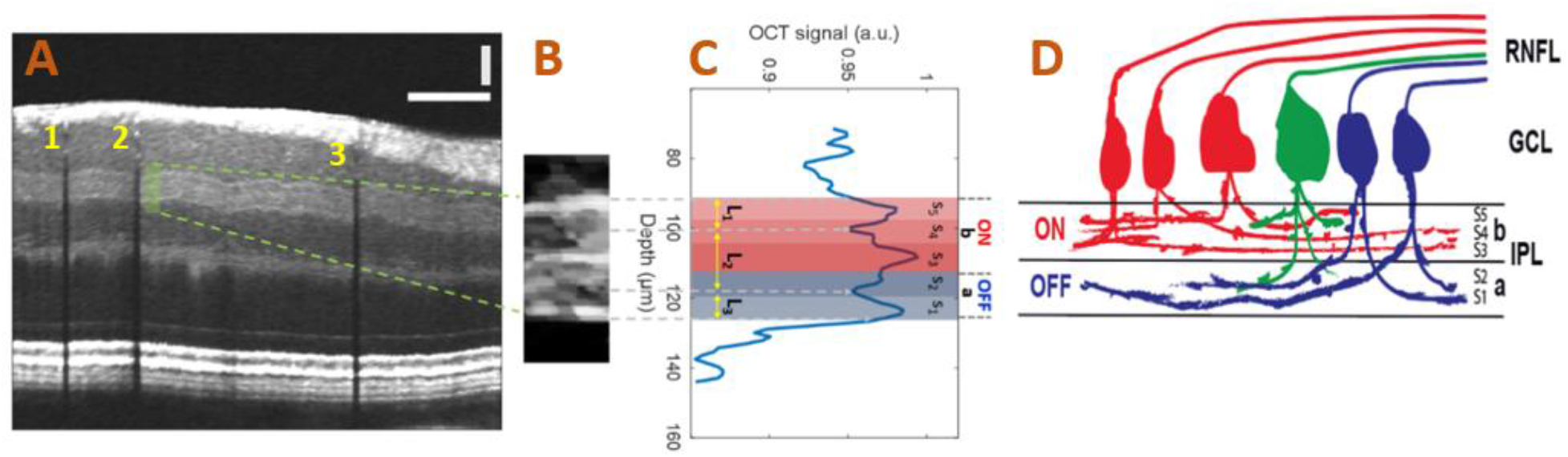
(a) A speckle-reduced vis-OCT image from a healthy subject. Horizontal bar: 500 μm; vertical bar: 50 μm; (b) Magnified view of the region highlighted by the dashed box in panel a; (c) Depth-resolved line profile of the IPL sublayers. We averaged 15 srA-lines, corresponding to approximately 88 μm along the lateral direction, within the highlighted region in panel a. (d) Illustration of the lamination of ganglion cells from RNFL to the IPL. The “red” ganglion cells (ON center) are laminating dendrites to the “b” sublamella of the IPL while “blue” cells (OFF center) laminating – to the “a” sublamella. The “green” ganglion cell is bi-laminating.

### Segmentation and thickness measurement of IPL sub-layers

With the improved vis-OCT imaging and post data processing method, we are able to distinguish the sublayer structures in the IPL. Specifically, we can detect three hyper-reflective bands in the IPL, with the top and bottom bands set the boundaries of the IPL. To measure variation in the fine lamella structure of the IPL, we identified three IPL sublayers that can be robustly measured from the averaged A-line profile with the following dividing lines (Figure 2c): (1) sublayer L_1_ measured from the top IPL boundary to a minimum of the first valley from the top of IPL; (2) sublayer L_2_ measured between minima of the first and second from the top of IPL valleys; (3) sublayer L_3_ measured from the minimum of the second valley from the top of IPL to the bottom IPL boundary. Therefore, L_1_ and L_3_ represent a part of the ON and OFF sublamina, respectively, while L_2_ includes both ON and OFF sublaminae. We measured the thicknesses of all the IPL sublayers manually from the averaged A-line profiles. The boundaries of the RNFL layer were also segmented manually at the same sampling locations as IPL sublayer measurements.

### Statistical analysis

Summary statistics were provided as mean and standard deviation (SD). Wilcoxon rank-sum test was used to compare the demographics between healthy and glaucomatous subjects. The coefficient of variations (CVs) of all three sublayers measured by vis-OCT as well as the entire IPL thickness was calculated to assess the intra- and inter-session repeatability. A linear mixed-effects model with a random intercept to account for intra-subject correlation was used to test whether parameters were different for glaucomatous and healthy subjects. Outcome measures included both the entire IPL thickness and its individual three sublayers. Statistical analysis was performed using R software version 3.5.2. A p-value of less than 0.05 was considered statistically significant.

## Results

Subject demographics are summarized in Table 1. Glaucoma subjects were older and showed lower MD and thinner RNFL than healthy subjects. To account for the age effect, we used a mixed-effect model for further comparisons. The global RNFL thickness was measured using a commercial NIR OCT system.

**Table 1.** Subject demographics

| subjects | Number | Age<br>(mean ± SD, years) | MD Range<br>(dB) | RNFL thickness<br>(mean ± SD, μm) |
| --- | --- | --- | --- | --- |
| Healthy | 9 | 45.4 ± 14.4 | -1.6 to 1.1 | 99.2 ± 16.2 |
| Glaucoma | 5 | 59.7 ± 13.4 | -26.4 to -7.7 | 59.6 ± 9.1 |
| P-value | - | 0.02 | 0.002 | 0.004 |

IPL sublayers were visible in all scans, both in glaucoma and healthy subjects. The imaging quality for healthy eyes was comparable to the glaucoma eyes without reaching statistical significance (p=0.07): an average QI for srB-scans used in the IPL layers thickness calculations was 65.4 ± 1.0 for healthy and 65.0 ± 0.8 glaucoma subjects.

### Intra-session repeatability

Intra-session repeatability results are summarized in Table 2. CVs showed good repeatability on all measured sublayers for both healthy and glaucomatous eyes. The variability of the entire IPL thickness measurements is significantly lower than the variability of the sublayer measurements for both healthy and glaucoma subjects. Among the sublayers, there was no significant difference in intra-session repeatability between glaucoma and healthy subjects.

**Table 2.** Intra-session repeatability for Healthy and Glaucoma Subjects

|  | Entire IPL (%) |  | Sublayer1 (%) |  | Sublayer 2 (%) |  | Sublayer 3 (%) |  |
| --- | --- | --- | --- | --- | --- | --- | --- | --- |
|  | median | range | median | range | median | range | median | range |
| Healthy (CV) | 3.1 | 0.0 - 4.3 | 5.6 | 5.1 - 6.2 | 6.9 | 3.5 - 8.8 | 5.6 | 5.4 - 5.6 |
| Glaucoma (CV) | 1.8 | 1.7 - 3.0 | 6.0 | 5.1 - 6.0 | 7.7 | 3.7 - 8.3 | 6.2 | 5.6 - 10.0 |
| P-value | 0.44 |  | 0.95 |  | 0.69 |  | 0.007 |  |

### Inter-session repeatability

Inter-session repeatability results are summarized in Table 3. CVs showed good repeatability on the entire IPL and the thickness of measured IPL sublayers in all healthy eyes (Table 3). The values were similar to intra-session study values.

**Table 3.** Inter-session repeatability for Healthy Subjects

|  | Entire IPL (%) |  | Sub layer1 (%) |  | Sublayer 2 (%) |  | Sublayer 3 (%) |  |
| --- | --- | --- | --- | --- | --- | --- | --- | --- |
|  | median | range | median | range | (median | range | median | range |
| Healthy (CV) | 2.9 | 1.6 - 4.3 | 5.4 | 5.2 - 5.5 | 6.8 | 5.1 - 7.8 | 5.6 | 5.1 - 6.2 |

### IPL sublayer thickness

IPL sublayer thickness results are summarized in Table 4. The entire IPL was significantly thinner in glaucomatous eyes than healthy eyes (p=0.003). The IPL sublayers L_2_ and L_3_ showed statistically significant differences between glaucoma and healthy subjects, while the sublayer L_1_ thickness did not. The sublayer L_2_ showed the greatest alterations in the glaucoma subjects. Age was not significant in the models.

**Table 4.** Measured IPL thickness and calculated p values based on *mixed-effects models* comparing the difference between Glaucoma and Healthy subjects

|  | Entire IPL | Sublayer 1 | Sublayer 2 | Sublayer 3 | RNFL |
| --- | --- | --- | --- | --- | --- |
| **Healthy**<br>(mean ± SD, $\mu\text{m}$) | 40.1 ± 1.7 | 11.4 ± 0.9 | 17.5 ± 1.4 | 11.3 ± 0.8 | 101.8 ± 13.8 |
| **Glaucoma** (mean<br>± SD, $\mu\text{m}$) | 36.2 ± 1.5 | 11.0 ± 0.9 | 14.2 ± 1.8 | 10.5 ± 0.8 | 59.8 ± 8.4 |
| **P Values** | 0.003 | 0.9 | 0.003 | 0.05 | 0.003 |

## Discussion

The integrative properties of RGC dendritic structure and function are critical for the visual function^9^. However, in prior studies using NIR OCT, the response of the IPL to glaucoma is considered insignificant compared with RNFL and combined GCL+IPL layers ^4 5^. In those studies, functional segregation of the ON and OFF pathways could not be investigated due to limited contrast and axial resolution. The newly developed vis-OCT has demonstrated the capability of revealing IPL lamination ^24, 30^, which makes it a unique technique to perform sublayer analysis on IPL *in vivo*. In fact, studies on RGC subtype variation in response to glaucoma in humans and animal models indicate that RGCs might exhibit different vulnerability to the disease insult ^34–43^. Quigley and colleagues had related greater conservation of RGC fibers with a smaller diameter compared with the RGC fibers with a larger diameter in a chronic glaucoma model of primates ^44^The parasol cells from glaucomatous eyes have smaller and less complex dendritic arbors, resulting in a significant reduction in total dendrite length and surface area ^45^. In mice, OFF-transient RGCs exhibited a more rapid decline in both structural and functional organizations, while ON-RGCs exhibited normal functional receptive field sizes ^46^. Duan et al. have shown that the α-RGC type was resilient to the ONC injury, while our studies suggested that a small subgroup of the ON α-RGC type was susceptible to the ONC ^16, 17^ The recent publication by Tran et al. ^47^ used the single-cell RNA sequence to create an atlas of RGC types and tracked type-specific RGC loss to the ONC injury ^47^ They described four different types of α-RGC, where two are resilient, and the other two are susceptible to injuries ^47^, which possibly explained the inconsistency among previous publications. In other words, there are dynamic changes in the IPL sublayers, which may offer us additional information on how subtype RGCs survive or die following the disease insults.

Here, our study took advantage of the novel imaging and analytic tools we recently developed to examine the dendritic changes in the IPL sublayers *in vivo*. For the first time, we were able to detect the subtle changes in the IPL sublayers. We have observed five bands in the IPL using vis-OCT imaging, which correlates well with the five IPL strata (Figure 2). Our results further showed that the reduction of the IPL in glaucoma is mainly due to the reduction in the thickness of the middle sublayer L_2_. It is of great interest to further match the histopathological changes in sublayers of the IPL with -the vis-OCT signal and investigate the underlying mechanisms in animal models of glaucoma. Our IPL sublayer analysis allows us to identify glaucoma phenotypes, which will also benefit personalized clinical care. For example, certain phenotypes may require aggressive treatment plans due to their potential rapid progression. Adjustments to the clinical visit schedule can be made accordingly.

Our results demonstrated the ability to produce repeatable thickness measurements of the IPL sublayers. To the best of our knowledge, this is the first attempt to quantify IPL sublayers of human eyes and compare the sublayer thickness between glaucoma and healthy subjects. High repeatability of IPL sublayer thickness, in both intra- and inter-session, makes the IPL sublayer analysis a strong candidate for a new biomarker for clinical glaucoma assessment. Moreover, investigating lamella-specific changes in the IPL in glaucoma may help elucidate the response of different types of RGCs to the disease.

Note that the median CV of the whole IPL thickness measurements was much lower than the median CV of the sublayer measurements in both healthy and glaucoma subjects. We attribute the lower variability of the entire IPL thickness due to the higher contrast between IPL and neighbor GCL and INL layers than between the IPL sublayers. Variability of the IPL sublayer L_2_ is the highest since it is the only sublayer without strong contrast at the boundary: the sublayer L_1_ and L_3_ have one boundary with a strong contrast: GCL/IPL for sublayer L_1_ and IPL/INL for sublayer L_3_.

While the entire IPL thickness was thinner in glaucoma than healthy eyes, the sublayer L_1_ (belongs to ON sublamella) did not show a significant difference between them. This suggests that the sublayer L_2_ (belongs to both ON and OFF sublamellae) and L_3_ (belongs to OFF sublamella) are likely responsible for the IPL thinning associated with glaucoma. However, observing the actual difference detected for both sublayers, the difference in sublayer L_3_ is within the optical axial resolution. Therefore, even with the statistical significance in the sublayer L_3_ difference, the primary measurable difference is attributed to the sublayer L_2_. This implies that glaucoma changes may take place in both ON and OFF lamellae, and due to observed changes in L_3_ but not in L_1_ - we can conclude that RGCs with dendrites stratifying in the OFF sublamella may be primarily altered in our glaucoma samples, who had moderate to advanced glaucoma.

The variability of the sampling locations to measure the IPL sublayer’s thickness was one of the limitations of the study. We aimed to measure IPL lamination from the same superior locations in the retinas measured from the reference srB-scan. However, the sampling location was dictated by the signal quality and patient’s motion. For example, in the srB-scan shown in Figure 2a, the IPL sublayers can be visually traced from the temporal to the nasal side. However, in some nasal locations, the IPL imaging quality was not sufficient for the quantitative measurements. Thus, lamination measurements in some nasal locations are not feasible.

Most measurements (69%) were taken from the srB-scan that is 2 and 3 superior to the srB-scan crossing the foveola. We took 31% measurement locations from 4 and 5 superior to the srB-scans. However, due to eye motions, we could not guarantee the exact location of the measurements. The uncertainty in the OCT imaging location due to eye motion in the range of 80 μm to 130 μm can be estimated from the data on the dispersion of the eye angle movement ^48–51^ using an effective focal length of the human eye as 16.7 mm ^52, 53^. Even with this relatively high subject to subject sampling location variability, both IPL and its sublayer measurement showed high repeatability, which suggests a possible homogeneity of the sublayer thickness in the parafoveal region.

## Conclusions

In this pilot study, we showed that speckle-reduction vis-OCT could provide a robust and repeatable quantitative assessment of the IPL lamination noninvasively in a small cohort of healthy and glaucoma volunteers. We visualized the five known morphological IPL strata with vis-OCT imaged IPL sublayers. We found that the majority of the changes in the IPL came from the thinning of the sublayer L_2_ in glaucomatous eyes.

